# Testing methods of linguistic homeland detection using synthetic data

**DOI:** 10.1101/2020.09.03.280826

**Authors:** Søren Wichmann, Taraka Rama

## Abstract

Two families of quantitative methods have been used to infer geographical homelands of language families: Bayesian phylogeography and the ‘diversity method’. Bayesian methods model how populations may have moved using a phylogenetic tree as a backbone, while the diversity method assumes that the geographical area where linguistic diversity is highest likely corresponds to the homeland. No systematic tests of the performances of the different methods in a linguistic context have so far been published. Here we carry out performance testing by simulating language families, including branching structures and word lists, along with speaker populations moving in space. We test six different methods: two versions of BayesTraits; the relaxed random walk model of BEAST 2; our own RevBayes implementations of a fixed rates and a variable rates random walk model; and the diversity method. As a result of the tests we propose a hierarchy of performance of the different methods. Factors such as geographical idiosyncrasies, incomplete sampling, tree imbalance, and small family sizes all have a negative impact on performance, but mostly across the board, the performance hierarchy generally being impervious to such factors.

## 1. Introduction

Information on the location of proto-languages is important in many studies of linguistic prehistory. Such origins may be of interest in and of itself, such as in the case of Indo-European, whose origins have received much attention and also been subject to controversy [1]. Language group origins may also, for instance, form the backbone for a study relating to linguistic typology or prehistorical areal linguistics [2]. When combined with information on dates for language groups [3], inferences about homelands become particularly powerful as contributions to world prehistory.

In the past, researchers had to rely mainly on *reconstructed* lexical items for clues to the origin of a language group. For instance, if a word for a particular biological species or material item which is diagnostic of a certain geographical or archeologically defined area, can be reconstructed for a proto-language, then the speakers of the proto-language can hypothetically be assigned to the area in question, e.g. [4]. In addition to this approach, known as linguistic paleontology, other types of evidence that can sometimes be drawn upon are old place names with known linguistic affiliations [5] or early loanwords showing different proto-languages to have been in contact [6]. These various approaches only promise to apply when a language group is already very well researched, and even then there are strong limitations since securely reconstructed and geographically diagnostic proto-words are hard to come by, and information on early loanwords and place names is most often not available.

As an alternative to linguistic paleontology, researchers have often inferred origins and directions of dispersion through an application of what can be called the center of gravity or diversity method. Apparently first introduced by Edward Sapir [7], the idea here is that the area of origin will usually see more diversity building up than areas in which members of a given language group were latecomers. For instance, Sapir himself argued [7] that the large Algonquian family of North America is more likely to have originated in the west than in the east because the most divergent languages are found in the west. The same type of argumentation has been applied to several families in South America [8], to Austronesian [9], to Sino-Tibetan (apparently) [10], and other families.

Recently, homeland detection methods of a more quantitative nature have appeared which only rely on a type of evidence which is available for any language group, namely basic *lexical items as attested in the extant languages*. The early part of the 2010’s saw both the application of biological (specifically Bayesian) phylogeographical approaches [11–14] and a quantitative implementation of the diversity approach [15]. These methods, however, have come with no warranty in terms of how well they perform on linguistic data and they have stirred some controversy in the linguistic community [16–17]. Regardless on one’s general view on quantitative methods, their results should be evaluated. But we rarely know the origin of a language group [15], so it is not obvious how to test the methods on empirical data. Even if we did collect information on cases of known origins and ran different methods to check their results we would not have enough data points to get good statistics on variability in performance or for getting insights about possible systematic causes for challenges to the methods.

The phylogeographic methods as applied to linguistic data are mainly based on the random walk models available in popular software such as BEAST 2 (henceforth simply ‘BEAST’) [18] and BayesTraits [19]. Apart from the random walk models, ancestral range reconstruction methods [20–22] can be used for estimating the ranges of internal nodes in a phylogeny. These ancestral range reconstruction methods are available as part of the RevBayes package [23]. They are useful for validating hypotheses regarding the probable geographical ranges of the ancestral nodes, given a set of candidate areas, but are not particularly suitable for inferring a set of probable geographical coordinates. For instance, the range estimation methods could be well-suited for validating a particular, geographical origin hypothesis [12] of a given language family such as Indo-European.

The lack of performance tests of linguistic homeland detection methods motivates this paper. The only other work of a similar nature that we are aware of is a recent manuscript still under review [24]. Even the biological literature is found wanting in comparative performance tests of phylogeographic methods, as has been repeatedly pointed out [24–26]. Tests have focused on technical aspects of individual methods, mainly as applied to epidemiology [27–28]. More work, congenial to our approach, has been done carrying out simulation-based tests of methods in the different, although related area of correlated evolution of continuous characters [29].

Phylogeography has grown into a diverse area—an excellent, recent survey [30] tabulates more than fifty combinations of softwares and methods—so we need to restrict our tests to the a representative sample, with attention to diversity and popularity of approaches.

We address the problem of lack of empirical testing data by producing synthetic data and then look at the performance of a range of methods in terms of the distance between the true and inferred homelands. The main set of simulated data consists of one randomly chosen language phylogeny and its dispersal in different real-world geographical settings under 1000 different origin scenarios. This dataset will represent the most important basis for our comparative assessment of the different methods. In addition, we provide some initial exploration of the effects of incomplete sampling by repeating tests for two pruned version of the tree, and, finally, preliminary assessments of the effects of the size of phylogenies and tree balancedness are carried out through the observation of the dispersal of 41 different-sized phylogenies on a plain lattice without geographical features. The next section describes the simulated data.

## 2. Data

We simulate a single language family having 20 languages. Each of the languages is associated with a 100-item word list containing words that have evolved over 120 time steps. The simulations are described in Online Appendix 3 to [31].^1^ Briefly, the ancestral language is constructed from an inventory of phonological symbols identical to the symbols used in the ASJP database [32]. Proto-words with semi-realistic shapes are constructed, and at each time step these words undergo phonological and lexical changes according to preset probabilities tuned to reality. Probabilities for speciation and extinction, also preset, allow lineages to grow into families of sizes that are not predictable but still probabilistically related to the parameters of the birth-death process and the number of time steps. The program was modified for the present paper to accommodate an output of 100-item rather than 40-item word lists and to include information about cognacy among the simulated words. It is found in the electronic supplementary material (SI-01) along with accompanying files and explanations. The program outputs five files, respectively containing the parameter settings, the topology, word lists for the terminal taxa in a tab-delimited format, the same word lists in the style of ASJP input files, and words lists where the words are shown in the shape they would assume if they did not undergo phonological changes. The information in this last file allows for easy extraction of cognacy relations.

Given that both information on cognate relations and actual forms having undergone phonological changes are provided, the simulated data could be used, for instance, to test techniques for automated cognate recognition, currently a rapidly developing field [33]. Here, however, the two alternative ways of presenting the lexical data serve to satisfy the requirements of the different methods that we are testing: the Bayesian methods that we test rely on information about cognacy—in general, characters are preferred over distances in Bayesian analyses because they allow for better estimates of uncertainty [34]—and are here supplied with the full, accurate information about the relations among 100 words; the diversity method, by the way it is constructed, relies on lexical distances and is here supplied with lists of 40 words in their actual shapes as input to the string distance computations. This somewhat less generous input is similar to real data that the method would use.

For the main part of the present study we selected one particular simulated family that seemed adequate in terms of its size—it is large enough to contain plenty of data for the methods to work with but not too large to be unwieldy. Its topology is shown in Figure 1. The number of time steps was set to 120, but the initial lineage happens to not split into other surviving lineages until step 77. Thus, the root of the lineage is anchored at step 76, which, for all practical purposes, can be considered step 0. The justification for pruning the initial branch is that the methods for detection homelands, even if successful, will only recover the most immediate location of the ancestral language. As is apparent from Figure 1, the two initial lineages split at the same time, 4 steps after the root. After a total of 44 steps (44 + 76 = 120) we reach the tips of the terminal branches. Since the simulations output word lists in the style of the input to a published, linguistic dating method [35], it is easy to translate time steps into calendar years, should one want a more palpable alternative. The family used here is 2297 years old when measured by lexical and phonological changes happening from the first split to the root. Thus, a time step, in this case, can be interpreted as representing 2297/44 ≈ 52 years.

**Fig. 1.**
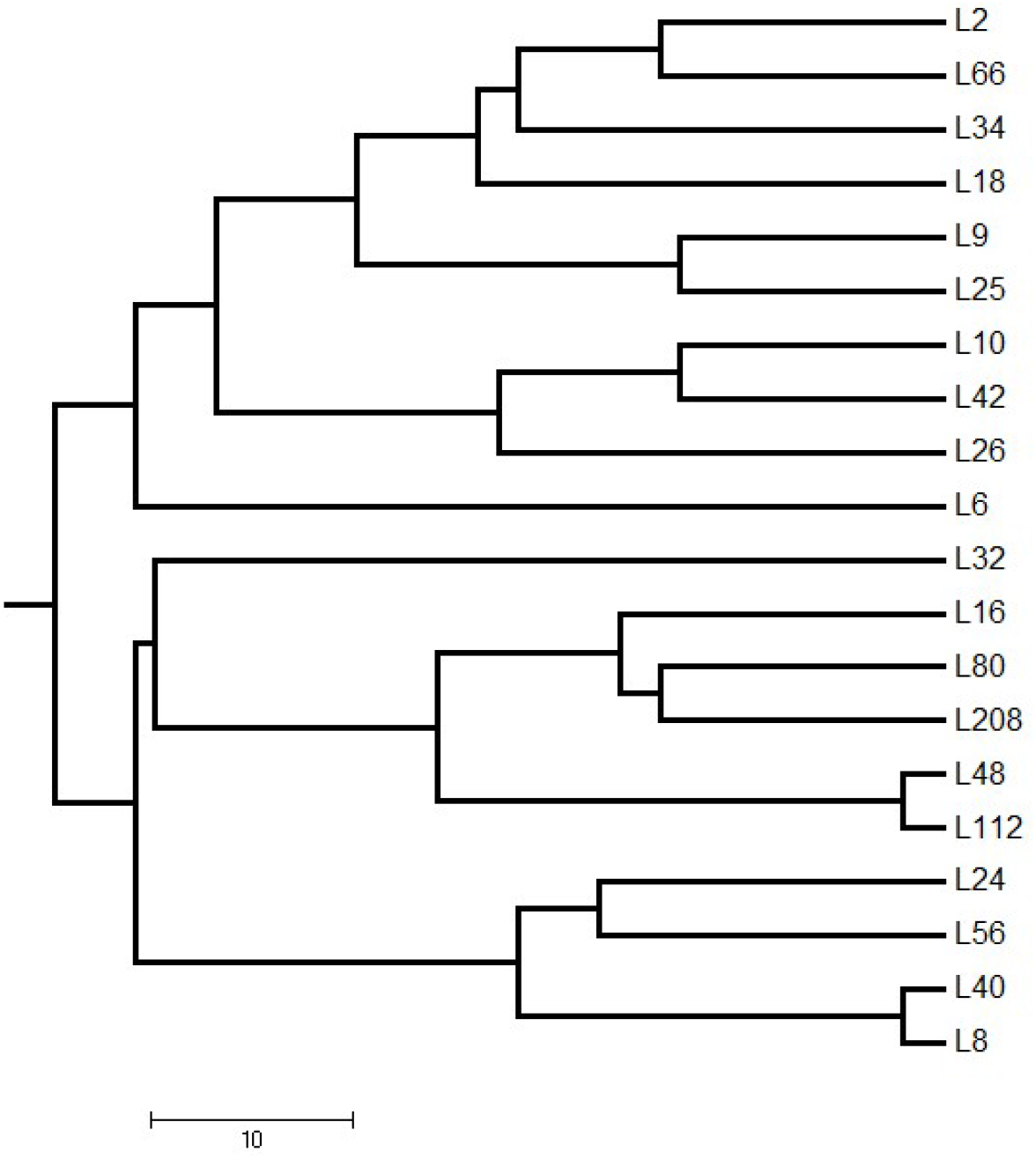
Simulated tree used in this study showing topology, branch lengths, and taxon names at the tip of each branch. The scale bar indicates time steps.

The structure of the tree does not translate directly into movements of languages. For instance, the fact that the two basal lineages have some time (4 time steps) to drift apart geographically does not mean that they have to—what happens in terms of movements depends on stochastic processes that we simulate separately. For simulating the geographical diffusion of languages we again use published software with a few modifications and make the updated software available here (electronic supplementary material SI-02, output in SI-03). Thus, for a full description and justification of the simulation procedure a paper [36] is already available.

The simulation process can be summarized as follows. Movements are constrained to any populated place on Earth, i.e. a place included in the geonames.org database. A starting point is found by randomly choosing from this set of populated places. At each time step there is a preset probability of moving to a new place within a square containing at least ch populated places. The ch parameter is here set to 800 since it was found to produce results that are realistic in terms of language densities [36]. Apart from the coordinates for the world’s populated places, the only input required is a tree structure such as the one displayed in Fig. 1, encoded as in the output for the language family simulation program.^2^ Using one and the same phylogeny, we produce 1000 diffusion scenarios. Since these play out in real-life geography, they will be sensitive to natural boundaries such as mountains, deserts, and bodies of water. If the diffusion starts on a small island containing less than 800 populated places there is a high probability that a lineage will make a jump away from the island; if it starts in a densely populated area the family may possibly never extend beyond this area. Places that are currently populated work as a proxy for the carrying capacity of different areas of the world. The kind of movement we simulate here may be called a semi-random walk, since it is a kind of random walk constrained to populated places. In spite of the constraint there is a possibility of jumps, so it is envisaged that a method for reconstructing a linguistic homeland will encounter both cases where languages have a relatively even distribution and cases where languages end up clustering in separate regions. Maps of all 1000 cases, showing the homeland, intermediate stations, locations of current languages, and inferred homelands similarly to Figure 2 below, as well as the script that produced the maps, are provided in the electronic supplementary material (SI-11).

**Fig. 2.**
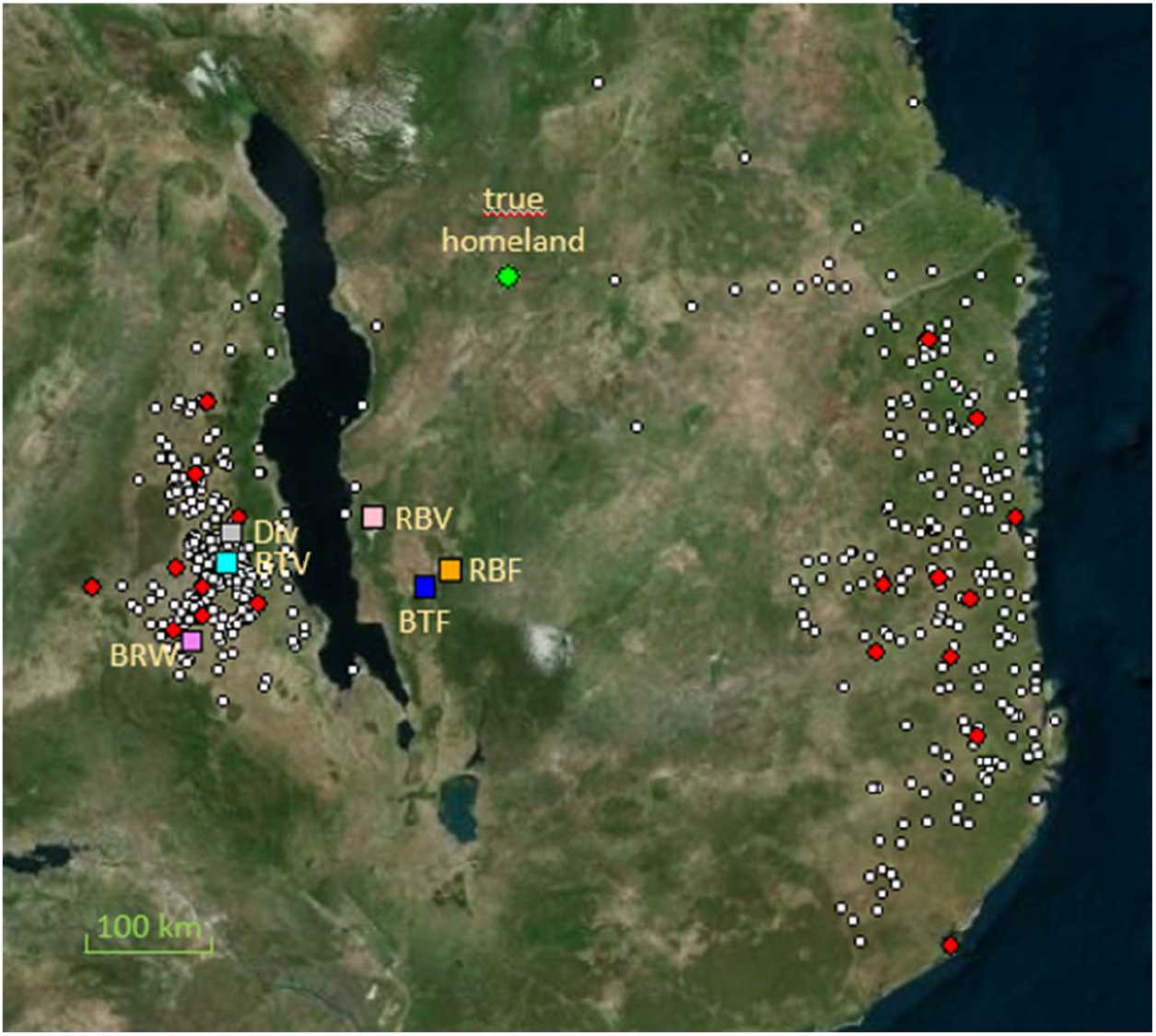
A simulation scenario where there were early jumps out of a homeland east of Lake Malawi (at 10.9 S, 36.15 E), causing all six methods to err. Red dots: extant languages; white dots: intermediate migratory stations; large green dot: true homeland; blue square: BayesTraits fixed rates (BTF); cyan square: BayesTraits variable rates (BTV); violet square: BEAST relaxed random walk (BRW); orange square: RevBayes fixed rates (RBF); pink square: RevBayes variable rates (RBV); gray square: Diversity (Div).

We choose to mainly work with a single simulated phylogeny unfolding in many different geographical locations because we hypothesize that the factor that will affect the performance of a homeland detection method the most, is the different geographical distributions of extant languages. Nevertheless, we also provide some initial exploration of other possible factors. *Incomplete sampling* is investigated by repeating experiments for two versions of a pruned tree. We refer to these versions as the ‘top’ and the ‘bottom’, with references to the visual display of Figure 1: the top contains the segment L2…L6, and the bottom the segment L32…L8. In other words, we check what happens when only one or the other half of the languages are available. In a more systematic investigation of the effect of incomplete sampling we should operate with a larger set of (incomplete) samples. Still, the preliminary investigation presented here will serve to check whether the variation in performance across methods is roughly similar under a condition of incomplete sampling. Moreover, we take a preliminary look at effects of *tree size* (number of languages) and *tree imbalance* by producing 41 different-sized phylogenies set to disperse from the center of a plain lattice—a symmetrical grid with no geographical features. Using a plain lattice for this experiment allows us to isolate the influence of factors other than geography. Other factors, such as change rates or extinction rates—factors known to affect ancestral state reconstructions [37]—could conceivably have been included in our investigation, but each additional parameter would imply a multiplication of computational efforts that are already large. We focus mainly on the role of the most interesting and complex parameter, namely geography, in the performance of homeland detection methods.

## 3. Methods

For each of the methods tested here and described in subsections (a)-(g), we provide all files needed for replication in transparently named folders contained in the electronic supplementary material (one folder per method).

### (a) Baselines (rand, centr, md)

As baselines we apply three approaches. The first approach, abbreviated ‘rand’, picks a random language in the family and assigns the homeland to the location of that language. Since this is merely a baseline and not a method meriting deeper investigation we just did a single run across the 1000 cases, even if the results would vary somewhat as different languages are selected. The second approach, abbreviated ‘centr’ for ‘centroid’ is built on the simple-minded assumption that the homeland is located in the center of the polygon represented by the extension of current languages. The centroid location is computed using the function ‘centroid’ of the R package geosphere [38]. In the third approach, abbreviated ‘md’ for ‘minimal distance’, we compute the average distance (as the crow flies) from each language to all the other languages. The location of the language that has the smallest average distance to the others is equated with the homeland.

### (b) Fixed rates model of BayesTraits (BTF)

The fixed rates model implemented in BayesTraits [19] is a Brownian motion model where the latitudes and longitudes are mapped to the three dimensional Cartesian coordinate system. The three dimensional coordinates are treated independently in this model. In the BTF model, there is a single parameter, namely the variance of the normal distribution, which is sampled using a Monte Carlo Markov Chain (MCMC) procedure along with the three dimensional coordinates. The fixed rates model takes a single tree with branch lengths and the geographical coordinates of the 20 languages as input and then reconstructs the internal nodes’ geographical locations using the MCMC procedure which is run for 1 million iterations.

We infer the phylogenetic tree of the 20 languages using a Metropolis-coupled MCMC (MC^3^) procedure. In the MCMC procedure, the cognate data of the 20 languages is supplied as an input to the MrBayes phylogenetic software [39]. We use the uniform model [40] that infers rooted trees using a binary continuous time Markov chain model of lexical evolution with sites weighted by a four-category discrete Gamma distribution. We perform two independent runs starting from two different randomly initiated starting points. Each independent run consists of three hot chains and one cold chain to navigate the multiple peaked tree landscape efficiently without getting stuck in local optima. An MCMC chain is run for 10 million iterations, where every 1000^th^ sample is written to a file. The first 25% of the iterations are discarded as burn-in. We constructed a majority consensus tree from the set of 7500 trees and used the majority rule consensus tree as input to both the BTF and the BTV BayesTraits models (subsection c), as well as to the RevBayes [23] model (described further below).

### (c) Variable rates model of BayesTraits (BTV)

The variable rates model is a relaxation of the single parameter Brownian motion [41], which assumes that the rate of change is fixed across all the branches. In this model, the branch lengths are allowed to shrink or expand reflecting large movements in space. In contrast to the MCMC models, where the number of parameters is fixed through the sampling process, the BTV model has two parameter changes: whether to scale a branch and to sample the scaling parameter of a particular branch. The decision to scale a branch increases the number of parameters by 1 and requires the use of a Reverse Jump MCMC (RJMCMC) procedure. The RJMCMC procedure does not easily converge, requiring long running times depending on the number of languages. In this paper, we run the model for 1 million iterations (sampled at every 1000th iteration) preceded by a burn-in of 100,000 iterations. In addition, we discard the first 500 samples and only use the sample with the best likelihood for evaluation purpose.

### (d) The relaxed random walk model of BEAST (BRW)

The relaxed random walk model as implemented in BEAST 2.6.3 [42] features a joint phylogenetic inference and phylogeographic model. In the phylogenetic inference model, lexical evolution follows a covarion model where some cognate sets are allowed to change faster than others. The tree model of evolution is based on a birth-death model where the height of the tree is drawn from a Gamma distribution with mean and standard deviation set to 1. The phylogenetic inference model also has a relaxed branch rates model where the branch rates are drawn from a uncorrelated lognormal distribution [43].^3^ The phylogeographic model is a relaxation of the Brownian motion model where the likelihood of latitude and longitude drawn from independent uniform distributions are estimated using a bivariate normal distribution. The parameters of the variance matrix of the bivariate normal distribution are scaled by a lognormally distributed scaler for each branch leading to a separate variance matrix for each branch. The model was run for 50 million iterations with every 1000th iteration written to a file. The first 50% of the samples were discarded as part of burn-in. Out of the remaining 25,000 samples, the geographical coordinates from the sample with the best posterior probability is used to evaluate the model.

### (e) RevBayes implementation of a fixed rates random walk model (RBF)

We implemented a simple model in RevBayes where the geographical coordinates are both drawn from uniform distributions of real numbers in the [−90, 90] range for latitudes and the [−180,180] range for longitudes. The phylogenetic tree is the majority consensus tree inferred from MrBayes software described in section 3(b). The standard deviation parameter of the normal distribution is assumed to be drawn from a uniform distribution on a log scale. The standard deviation parameter along with latitudes and longitudes are sampled using an MCMC chain. As part of burn-in, the chain was run for 5000 iterations followed by a run for 50,000 iterations sampled at every 100th iteration to reduce autocorrelation.

### (f) RevBayes implementation of a variable rates random walk model (RBV)

The RBV model [44] allows for variable rates of evolution among branches using a relaxed model where each branch’s rate is allowed to shift or not to shift. If there is no shift, the rate for the branch is the same as the rate for its parent branch. If there is a rate shift, the parent branch’s rate is scaled by a scalar that is sampled through an MCMC procedure. The rate parameter at the root node is sampled through an MCMC move. The rate shift multiplier for a branch is drawn from a Gamma distribution with mean 1 and variance set to 0.33. The prior probability that there is a rate shift is given as the expected number of rate shifts (5) divided by the number of branches (20 – 1 = 19). The rate shift multiplier being 1 (i.e., no rate shift) or not 1 (drawn from the above mentioned Gamma distribution with mean 1 and variance 0.33) is sampled using a Reverse Jump MCMC move since a value not equal to 1 would increase the number of parameters in the model by 1. We ran the model for 100,000 iterations with every 200th iteration written to file. The run was preceded by a burn-in of 5000 iterations. RBV uses the same phylogenetic tree and nexus files as the BTF (see Section 3b) model as input.

### (g) The diversity method (Div)

This method represents a quantitative implementation of the old idea in linguistics and biology [7,45] that the area of greatest diversity of a family/species is most likely the homeland. We follow [15], where the method is presented and results for empirical data from across the world’s language families are discussed. An actual program for doing the calculations is published for the first time in the electronic supplementary material (SI-10). Since the method is already amply described [15], we only provide a summary here. A basic tenet of the method is that the location of one of the current languages can (approximately) be identified with that of the homeland.

Given this assumption, the question becomes how to assign diversity values to different languages. This question is answered by assuming that larger linguistic distances coupled with greater proximity to the linguistically distant relatives means more diversity. This solution is then implemented quantitatively by computing all pairwise linguistic distances as well as all pairwise geographic distances (as the crow flies) among the languages. The linguistic distances are average string distances across word lists, the particular string distance used being defined as LND or Levenshtein Distance Normalized—the Levenshtein distance divided by the length of the longest of the two words compared—divided by the mean LDN across word pairs not referring to the same concept. This is called the LDND (Levenshtein Distance Normalized Divided) [46]. Now a diversity index DL for each language L is calculated as the mean of the proportion of linguistic to geographical distance between a given language and each of the other languages. The location of the language with the highest DL value is assumed to be the homeland.

## 4. Results

We first look at results for the different methods as distances in km (as the crow flies) from the true to the inferred homelands. For the Bayesian methods we initially ignore the fact that they provide a set of plausible homelands, i.e., inferred areas rather than single locations—for the purposes of a quantitative performance comparison it is necessary to operate with a single location whose validity can be assessed as a number. Nevertheless, later in this section we also take into account the full set of credible origins (95% Highest Posterior Density or HPD intervals) in a comparison among the Bayesian methods.

For the purpose of identifying the ‘best’ homeland according to the Bayesian methods we proceed as follows. For BayesTraits (BTF, BTV) we select the location associated with the highest likelihood (the software only outputs likelihoods, not posterior probabilities) and for BEAST (BRW) and RevBayes (RBF, RBV) we select the location with the highest posterior probability.

Table 1 provides mean and median absolute errors (distances) (for detail on individual cases see the file summary_errors.txt in the electronic supplementary material, SI-11). Across the table, we observe large differences in means and medians, suggesting that errors are not normally distributed, outliers having a large impact. While the absolute errors are suggestive of some performance differences they should be taken with a grain of salt since they are generally highly correlated with the mobility of a family. The latter can be measured as the total distance traversed at all time steps by the twenty lineages. For the centr (centroid) baseline the correlation is small although still significant (*r* = .173), but this is also the ‘odd man out’ in terms of performance. For the other baselines and methods the correlations, which are all significant at the *p* < .00001 level, are in the range .657 ≤ *r* ≤ .721. Given the correlations between absolute error and mobility we prefer to assess performance using relative error, calculated as the absolute error divided by total distance traversed. These numbers are also displayed in Table 1, multiplied by 100,000 and rounded to integers for easier viewing (see summary_relative_errors.txt in the online electronic supplementary material, SI-11, for more detail).

**Table 1.** Mean and median absolute errors (in km, rounded) and mean and median relative errors. Abbreviations: rand = random baseline; centr: centroid baseline; me: minimal distance baseline; BTF: the fixed rates model of BayesTraits; BTV: the variable rates model of BayesTraits; BRW: the random walk model of BEAST; RBF: the fixed rates model of RevBayes; RBV: the variable rates model of RevBayes; Div: the diversity method.

| Error | rand | centr | md | BTF | BTV | BRW | RBF | RBV | Div |
| --- | --- | --- | --- | --- | --- | --- | --- | --- | --- |
| Mean abs. err. | 214 | 353 | 153 | 148 | 172 | 179 | 167 | 172 | 183 |
| Med. abs. err. | 156 | 160 | 106 | 103 | 114 | 123 | 118 | 125 | 131 |
| Mean rel. err. | 1137 | 2051 | 794 | 781 | 884 | 931 | 878 | 915 | 955 |
| Med. rel. err. | 1008 | 997 | 682 | 679 | 749 | 805 | 773 | 791 | 833 |

The results for mean relative errors in Table 1 allow us to propose a hierarchy of methods and baselines, as follows:

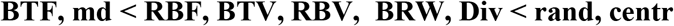

To construct this hierarchy, we first ordered the methods and baselines, separating them by the less-than symbol. Subsequently we applied a pairwise Wilcoxon rank sum significance test to assess the differences between *neighbors* in the hierarchy and replaced the smaller-than sign by a comma when neighbors were not significantly different at the .05 level. This step caused BTF/md, RBV/BRW/Div, and rand/centr to merge. We then iterated this procedure, also replacing the less-than sign with a comma when there were non-significant differences between members of neighboring *groups*. The resulting picture is the one where there are just three groups. BayesTraits (specifically BTF) works best, followed by RevBayes (RBF, RBV), BRW, and Div, which are rather indistinguishable. All the methods beat the baselines rand and centr, which provides a good sanity check on the results. A surprising result, however, is that the md (minimal distance) baseline shares the winning position with BayesTraits. This baseline does not draw on any linguistic information, only geographical coordinates. It is ‘one half’ of the diversity method (Div) in the sense that Div uses average geographical distances in its denominator in the calculation of diversity indices used to select the best homeland language.

A pairwise correlation of relative errors across methods, also including the baselines, shows that these errors are highly correlated, except for the baselines rand and centr. For all pairs of methods (also including md) correlations lie in the range .809 < *r* < .935 (see the electronic supplementary material, SI-11, for more detail). Given the overall relative similarity of performances it does not seem productive to look for factors that systematically incur *differences*. Instead, we will look for factors that affect the performance in *similar ways* across methods. The large differences in mean and median relative errors (Table 1) indicate that the variability of performance is mainly due to large outliers. Inspection of pictures showing inferred and true homelands for each method (included in the electronic online supplementary material folder SI-11) suggest that outliers are mainly due to a situation where all current languages are found in areas other than that of the homeland, presumably due to early geographical movements pertaining to the two main lineages. Figure 2 shows such a situation where languages have ended up in two different areas, apparently as a result of early jumps out of the homeland of both major lineages. All the six methods infer wrong homelands, although there are some differences in the magnitude of the errors.

If it is correct that large outliers in errors are mainly due to early jumps out of the homeland, then we can gather from the tree structure in Figure 1 that these jumps would have occurred during the first 4 time steps of the simulations. In order to corroborate this hypothesis we first compute ‘early jump indices’ as the percentage of the distances traversed during the first 4 time steps out of the sum of all distances traversed (results are in the file percent_early.txt in the online supporting material folder SI-11). We then gauge the impact of early jumps on errors in two different ways, as follows: (a) early jump indices are correlated with relative errors across the 1000 cases; (b) following the traditional way of defining outliers in a histogram, taking a larger outlier to exceed a value of 1.5 times the interquartile range, we identify outliers in the early jump indices as well as in the relative errors and then compute an agreement index as the percentage of large error outliers that pertain to cases which are also large outliers in terms of their early jump indices. If errors are in a large part due to early jumps, the correlations produced in approach (a) should be high, but the correlations are based on all 1000 sets, so factors other than early jumps affect the results. Approach (b) is more focused just on the early jumps. Results for all baselines and methods except for centr, for which the correlations (approach a) were non-significant, are displayed in Table 2.

**Table 2.** The relationship between the presence of early jumps and absolute errors. The row ‘correlation’ shows correlations between the percentage of the distances traversed during the first 4 time steps out of the sum of all distances traversed and relative error. The row ‘agreement’ shows the percentage of large error outliers that pertain to cases which are also large outliers in terms of their early jump indices. Abbreviations are the same as in Table 1.

| Approach | rand | md | BTF | BTV | BRW | RBF | RBV | Div |
| --- | --- | --- | --- | --- | --- | --- | --- | --- |
| correlation | .418 | .681 | .655 | .632 | .604 | .569 | .568 | .545 |
| agreement | 44.4 | 71.4 | 69.4 | 64.5 | 55.6 | 56.4 | 60.6 | 46.7 |

Table 2 shows that the two approaches to measuring the error rates’ sensitivity to early jumps agree (Spearman’s *ρ* = .905) and that they both indicate relatively high sensitivities. Thus, early jumps are a problem for all methods.

It is interesting to see how much the mean errors would reduce if we could identify cases of early jumps and exclude them. By the traditional criterion just mentioned there are 60 such outliers. Leaving them out does not change much for the centroid and random baselines— respectively a reduction of 1.9% and 5.2% of the mean error. For the other methods and baselines the reduction lies in the range 6.9%-9.2%. If the 60 cases are excluded, the above performance hierarchy stays the same apart from a local trade of places between BTV and RBF, two methods whose performance was not significantly different anyway. In conclusion, early jumps are not the sole cause of error, even if they pose a problem across the board, and they do not explain why different methods have different performances.

The only other simulation-based study looking at the performance of Bayesian phylogeography (but excluding other approaches from its purview) [24] similarly finds directed, migratory steps to be problematical. In this study simulations were specifically designed to assess the performance of the relaxed random walk and constant directional random walk models as implemented in BEAST 1.10.4 in a situation where languages have spread in a directional fashion versus a situation more akin to a random walk. The authors find that both models produce large errors when the language spread is directional.

Up to this point we have used as a criterion of performance the distance between the true homeland and a single, ‘best-candidate’, homeland even for the Bayesian methods, although these methods actually produce a range of credible candidate locations rather than a single location. This was done in the interest of comparability. Now we look at whether conclusions might look different when considering the 95% HPD (highest posterior density) intervals, extracting these as done in [12]. These intervals correspond to geographical ‘swarms’ of dots that strongly tend to form circular or elliptical shapes. Therefore, one can assess a result by fitting a convex hull to the swarm and determine whether the true homeland lies inside or outside of this fitted periphery. This test, however, is confounded by the fact that the area of the convex hull can vary tremendously. For instance, if one method proposes an area almost corresponding to the extension of the entire family and the true homeland falls *within* this area, is it then fair to say that the result is better than the result of some other method whose output is an area which is ten times smaller and lies in the vicinity of the true homeland but does not contain it? In order to take this issue into account we record not only hits and misses, i.e., containment within vs. absence of the true homeland from the fitted convex hull, but also measure the area of the latter. This allows us to produce a performance measure where the number of hits out of the 1000 cases is weighted (divided) by the mean size of the areas of the convex hull. Results for the five Bayesian methods are displayed in Table 3. While BRW has the highest absolute hit rate, the weighted hit rate is the lowest. If the weighted hit rate were to be used as a performance measure, the internal ranking of the Bayesian methods would correspond to what we saw earlier, except that BTV and RBF, whose ranking was actually not significantly different, would swap ranks.

**Table 3.** Number of hits (location of true homeland within a convex hull fitted to coordinates pertaining the 95% HPD intervals), area (in km2) of the convex hull, and the ratio between the two.

|  | BTF | BTV | BRW | RBF | RBV |
| --- | --- | --- | --- | --- | --- |
| Hits | 340 | 292 | 789 | 609 | 622 |
| Mean Area | 17,871 | 30,958 | 232,715 | 122,856 | 129,173 |
| Ratio | 0.01902 | 0.00943 | 0.00339 | 0.00496 | 0.00482 |

In defense of BRW it can potentially be argued that it is not in the spirit of Bayesian inference to disqualify a method because of the generosity of its credible interval. Nevertheless, if this interval is so huge as the case is here—the mean area being 13 times as large as that of BTF—*and* when it is still completely misspecified in more than 20% of the cases, then the method would simply cease to have much practical use. We conclude that the performance hierarchy already proposed is not affected when taking 95% HPD intervals into account.

We now go on to looking whether the hierarchy is affected by sample size, tree size, and tree imbalance. Since it now seems justified to use the distance between the true homeland and a single, ‘best-candidate’ inferred homeland, this is going to be our sole performance measure for the rest of the section.

For the experiments where only one half of the languages (either the ‘top’ or the ‘bottom’ segment of Figure 1) is made available for inferring homelands, the mean and median relative errors are as indicated in Table 4. Compared to the relative errors using the full tree (Table 1), we see numbers that are around twice as large when only one half of the languages (whether one or the other half) is used to infer the homeland. The error increments are similar across methods.

**Table 4.** Mean and median relative errors (in km, rounded) for the pruned tree.

| Version | Error | rand | centr | md | BTF | BTV | BRW | RBF | RBV | Div |
| --- | --- | --- | --- | --- | --- | --- | --- | --- | --- | --- |
| ‘Top’ | Mean | 2128 | 3468 | 1737 | 1736 | 1805 | 1906 | 1747 | 1814 | 1937 |
|  | Median | 1820 | 1875 | 1438 | 1441 | 1478 | 1604 | 1500 | 1531 | 1670 |
| ‘Bottom’ | Mean | 2521 | 3629 | 2006 | 1901 | 2037 | 2071 | 1951 | 2006 | 2220 |
|  | Median | 2276 | 2098 | 1711 | 1659 | 1750 | 1796 | 1645 | 1670 | 1933 |

Indeed, a hierarchy based on the ‘top’ sample would be exactly the same, and a hierarchy based on the ‘bottom’ sample would differ only in local swaps in the ranks of md/RBF and RBF/BTV. (For more detailed results and all relevant files see SI-12).

A final additional experiment was carried out in order to preliminarily gauge the impact of tree size and tree imbalance. We produced a large set of simulated phylogenies and sampled 41 trees with sizes spread across the range from 4 to 45 languages. In this experiment, all the Bayesian methods were supplied with true topologies as starting trees in order to further control the comparisons. (For more detail see SI-13). The mean weighted imbalance, *I_w_* [47] was computed using the fusco.test() function of the R package caper [48] after transforming the encoding of tree structure output by the simulations into newick format. We did not attempt to artificially induce different degrees of imbalance, but simply observed the imbalance of the different trees produced (mean *I_w_* = .572, minimum *I_w_*: 0.3, maximum *I_w_*: 1). In Table 5 we report mean and median relative errors in the same style as for the full and pruned 20-language tree above.

**Table 5.** Mean and median relative errors, variable-size phylogenies on a lattice.

| Error | rand | centr | md | BTF | BTV | BRW | RBF | RBV | Div |
| --- | --- | --- | --- | --- | --- | --- | --- | --- | --- |
| Mean | 1265 | 1276 | 937 | 762 | 874 | 1200 | 797 | 917 | 1030 |
| Median | 928 | 754 | 452 | 467 | 594 | 802 | 482 | 696 | 569 |

A performance hierarchy extracted from the mean errors displayed in Table 5 would be the same as for the 20-language phylogeny in real-world geography, except that the minimal distance (md) baseline would move down to the fifth position in the hierarchy. These results merely serve to *suggest* that the performance hierarchy is impervious to tree size. As it turns out, differences in means are nearly all non-significant by a Wilcoxon test (the exceptions nearly all pertaining to the extreme case of the random baseline), so we would need a larger sample for full confirmation.

As for the effects of tree size and imbalance, these are displayed in Table 6. All methods and baselines exhibit small, negative correlations between tree size and *relative* error, meaning that they work somewhat better the larger the trees are. A study of ancestral state reconstruction [37] similarly found that accuracy grows with tree size.

**Table 6.** Correlations (Pearson’s *r* and associated *p*-values) between relative errors and each of the variables tree size (*N*) and mean tree imbalance (*I_w_*) for the variable-size phylogenies on a lattice.

| Method | $N$ | | mean $I_w$ | |
| --- | --- | --- | --- | --- |
| | $r$ | $p$ | $r$ | $p$ |
| BTF | -0.581 | 0.00007 | 0.309 | 0.04923 |
| BTV | -0.420 | 0.00621 | -0.144 | 0.36810 |
| BRW | -0.628 | 0.00001 | 0.357 | 0.02202 |
| RBF | -0.536 | 0.00031 | 0.246 | 0.12100 |
| RBV | -0.625 | 0.00002 | 0.488 | 0.00121 |
| div | -0.467 | 0.00208 | 0.089 | 0.57900 |
| random | -0.558 | 0.00015 | 0.253 | 0.11110 |
| centroid | -0.325 | 0.03788 | 0.141 | 0.38090 |
| mindist | -0.538 | 0.00029 | 0.295 | 0.06091 |

Tree imbalance shows small, positive correlations with relative error for the fixed rates model of BayesTraits (BTF), the relaxed random walk model of BEAST (BRW), and the variable rates model of RevBayes. Thus, for these methods, more imbalance tends to lead to a greater error. We are not certain how to interpret the absence of significant positive correlations for the variable rates model of BayesTraits (BTV) and the fixed rates model of RevBayes. It may well be that the sample is simply too small to produce *p*-values even at that *p* < 0.05 level. The absence of correlations for the diversity method, which is not tree based, and the baselines, which are also not based on trees, is completely expected, however.

## 5. Discussion

In this paper, we have tested different linguistic homeland detection methods comparatively in 1000 different scenarios and, more preliminarily, under different conditions of incomplete sampling and variable tree sizes. The result was a robust performance hierarchy.

Some comments on qualitative aspects of performance are also in order. In terms of runtime, the diversity method (Div) and the minimal distance (md) baseline require less than a minute on a laptop while the relaxed random walk model of BEAST (BRW) requires close to an hour for each case of phylogeny of 20 languages on a server, with the other methods being intermediary. The Bayesian methods require cognate information, while the diversity method only requires word lists, and using the minimal distance does not even require word lists. Div and md will avoid assigning homelands to uninhabitable areas, including water bodies, since they are restricted to populated locations. All the Bayesian methods give samples of coordinate sets associated with posteriors measuring confidence, as opposed to Div and md, which just output a single location. Such qualitative factors can feed into the choice of a particular method. In the case of BEAST, the computing time may be prohibitive in a study involving very large or numerous language groups. Typically, cognate information may not be accessible for all the language groups even though word lists are available. This study suggests that BTF may be the preferred method when the cognate information required to infer a phylogenetic tree is available (such information may be produced manually or through an automated procedure [49]). When it is not, Div can be used, and its performance is not going to be radically inferior in spite of the more limited information that it draws upon. The surprisingly effective md baseline may serve as a quick way of generating a hypothesis about the location of a homeland, and it is the only possible approach in cases where no linguistic information is available.

As for the interpretation of our quantitative results, the question naturally arises why one method seems to work better than another. Here we concentrate on the Bayesian methods. We found that the fixed rates variant of Brownian motion as implemented in BayesTraits (BTF) works better than the Brownian motion models implemented in RevBayes (RBF, RBV) and BEAST (BRW). We think that the main reason is that the BayesTraits variant transforms the geographical coordinates into three dimensional Cartesian coordinates, which is more suitable for modeling geographical movements in terms of Brownian motion than the RevBayes or BEAST variants, which assume a rectangular flat earth surface for modeling geographical expansion. As for the reason why the variable rates models as implemented in BayesTraits and RevBayes do not work as well as the fixed rates BayesTraits model, this may relate to the fact that the simulations have a probabilistically fixed lexical replacement rate that is not related to the branch length. BEAST ends up in the lower end of the performance hierarchy. This may be due to model overparameterization: BEAST’s relaxed random walk model allows for each branch to have a separate rate, and there may simply not be information to estimate each branch-specific rate as a free parameter [44]. This does not happen in the variable rates models in BTV and RBV. Another possible reason for the low performance of BEAST could be the joint inference of both phylogeny and homelands. Overparameterization and the joint inference procedure could both contribute to explaining not only the high relative errors but also the large HPD areas.

Throughout this paper we have not always kept a straight conceptual distinction between softwares and methods, but it is probably true to say that one cannot expect the same output from any two of the softwares used here even when settings are the same, and much of the mechanics under the hood is sparsely documented. Thus, there are differences residing in softwares, but our main focus has been on methods. If we were to focus on the comparison of softwares we could have designed the study differently, for instance by fixing the tree topology for BEAST for a more direct comparison with BayesTraits and RevBayes, or, reversely, RevBayes could have been set up to simultaneously infer phylogeny and homeland. But, as mentioned in the Introduction, there are more than fifty combinations of methods and softwares available [30], so we had to be selective (cf. [29] for a similarly selective approach in the area of tests of methods in correlated evolution).

In the future, alternative simulation approaches may contribute to supporting or nuancing these results, and it would also be interesting to see studies directly aimed at testing the tentative explanations given here for the variation in performance. Of particular interest here are performance effects of a variable vs. fixed rates model and of a joint vs. separate inference of phylogenetic structure and geographical movement. Meanwhile, the field of phylogeography is developing rapidly, making it a moving target when it comes to testing. Currently the trend seems to be towards models that are richer in relevant biological [50] and environmental [51] information. Such enriched models may perform better, but we also suspect that they will pose new challenges for comparative testing.

## 6. Conclusion

The practical consequences of our main results of testing different methods and baselines for inferring the geographical origins of language groups can be summarized as follows. When cognate information is available, the fixed rates model of BayesTraits (BTF) seems preferable, although we cannot exclude that the variable rates model of BayesTraits (BTV) may be more suitable when it comes to real-life data. When cognate information is not available, the diversity method (Div), which requires neither a tree nor cognate information, may be applied, and its results will not be radically different from methods implemented in BayesTraits and largely indistinguishable in quality from those of RevBayes and BEAST. The minimal distance (md) baseline may have a real utility as a quick approximation to a hypothetical homeland, and is useful especially when only language locations are known.

## Acknowledgments

SW’s research was carried out under the auspices of the project “The Dictionary/Grammar Reading Machine: Computational Tools for Accessing the World’s Linguistic Heritage” (NWO proj. no. 335-54-102) within the European JPI Cultural Heritage and Global Change programme (http://jpi-ch.eu/). It was additionally funded by a subsidy from the Russian government to support the Programme of Competitive Development of Kazan Federal University and a major project from National Social Science Fund of China (no. 19ZDA300).

## Supplementary material

Electronic supplementary material is available online at https://figshare.com/articles/dataset/Testing_methods_of_linguistic_homeland_detection_SI/133 08926. It contains all programs and in- and output files used in this paper as well as explanatory notes.

## Footnotes

1 Available at https://datadryad.org/stash/dataset/doi:10.5061/dryad.4gg07.

2 The matrix-style tree encoding format, which we refer to as the ambar format, after a restaurant in Kazan called Staryj Ambar, where some aspects of it was developed, can be transformed to the more well-known newick format (also named after a restaurant) using the script ambar2newick.R provided in SI-01.

3 The XML file was crafted based on the tutorial at https://taming-the-beast.org/tutorials/LanguagePhylogenies/.

## Notes

https://doi.org/10.6084/m9.figshare.13235093

### Competing Interest Statement

The authors have declared no competing interest.

### Summary of Updates

This version has more extensive references to relevant literature, but otherwise no major changes.

